# Drift as a driver of language change: An artificial language experiment

**DOI:** 10.1101/2021.03.26.437270

**Authors:** Rafael Ventura, Joshua B. Plotkin, Gareth Roberts

## Abstract

Over half a century ago, George Zipf observed that more frequent words tend to be older. Corpus studies since then have confirmed this pattern, with more frequent words being replaced and regularized less often than less frequent words. Two main hypotheses have been proposed to explain this: that frequent words change less because selection against innovation is stronger at higher frequencies, or that they change less because stochastic drift is stronger at lower frequencies. Here, we report the first experimental test of these hypotheses. Participants were tasked with learning a miniature language consisting of two nouns and two plural markers. Nouns occurred at different frequencies and were subjected to treatments that varied drift and selection. Using a model that accounts for participant heterogeneity, we measured the rate of noun regularization, the strength of selection, and the strength of drift in participant responses. Results suggest that drift alone is sufficient to generate the elevated rate of regularization we observed in low-frequency nouns, adding to a growing body of evidence that drift may be a major driver of language change.

## Introduction

George Zipf noted a series of statistical regularities in natural languages (Zipf, 1949). Best known among these is the power law linking word frequency and rank (Piantadosi, 2014; Moreno-Sanchez et al., 2016). Another well-known regularity is the negative correlation between word frequency and length (Kanwal et al., 2017; Mahowald et al., 2013; Piantadosi et al., 2011; Sigurd et al., 2004). Indeed, similar patterns have been found in some forms of animal communication (McCowan et al., 1999, 2005; Suzuki et al., 2005). But Zipf also made the less well-known observation that “the less frequent words contain an increasing proportion of the later adoptions” (Zipf, 1949, p.12), meaning that rare words are more likely to be recent borrowings or coinages. Recent studies have confirmed that frequent words have lower rates of replacement and regularization (Pagel et al., 2007; Lieberman et al., 2007; Gray et al., 2018).

However, it is not clear why frequency of use should predict rates of lexical regularization and replacement. Zipf hypothesized that this pattern results from a trade-off between a pressure for successful communication and a pressure for efficiency (Zipf, 1949). That is, words that occur more frequently serve a greater communicative need and are thus under stronger pressure not to be replaced or regularized. In cultural evolutionary terms, this is to say that the pattern is driven by selection. Selection in this context is any *directional* bias in the acquisition, processing, or production of language (e.g., a preference for one form over alternative forms for the same meaning). Several studies have found evidence for selection in language change (Amato et al., 2018; Sindi and Dale, 2016; Stadler et al., 2016). Selection could be responsible for the lower rate of regularization among high-frequency words if selection against innovations is stronger during acquisition, recall, or production of high-frequency terms (Pagel et al., 2007).

Although many social, cognitive, and linguistic factors can give rise to selection in this sense, an important and simple source of selection is relative frequency during language learning. When two alternative forms for the same meaning occur at different relative frequencies, both child and adult language learners tend to regularize by eliminating the less frequent of the two forms (Hudson Kam and Newport, 2005; Hudson Kam and Newport, 2009; Reali and Griffiths, 2009; Smith and Wonnacott, 2010; Smith et al., 2017). Relative frequency may also interact with and bolster the effects of other linguistic or social sources of selection (Labov, 2001). In other words, language learners may favor the more frequent of alternative competing forms, giving rise to selection for the more frequent form and against the less frequent one. Thus, if biases of this kind are stronger for more frequent words, then regularization will be lower among high-frequency words, meaning that the negative correlation between word frequency and regularization could be driven by selection.

But another possibility is that the pattern is simply driven by drift, with infrequent words regularizing and being replaced more by chance because sampling variance is greater at lower frequencies (Reali and Griffiths, 2010; Newberry et al., 2017). By “drift” we mean any source of unbiased stochasticity, or sampling error, in the acquisition, processing, or production of language. Several studies have detected signatures of drift in language change (Bentley, 2008; Hahn and Bentley, 2003; Newberry et al., 2017). A similar mechanism could explain the higher rate of regularization among low-frequency words. Drift is stronger at lower frequencies as a consequence of the statistical fact that sampling error is stronger in smaller samples. As language learners sample a finite set of language-related stimuli, variance in the frequency of alternative forms that language learners encounter is greater for lower-frequency words. As a result, language learners may be more likely to acquire one form at the expense of another when words occur at low frequencies. For example, if an English speaker were to choose past tense forms in proportion to how often they encountered each form during learning, then one might win out simply because chance exposed them to that form more often than the other. Because of the relationship between frequency and sampling error, regularization and replacement may thus occur to a greater extent in less frequent words as a result of drift.

Both hypotheses are plausible, and the few studies on this question have been inconclusive (Pagel et al., 2007; Lieberman et al., 2007; Newberry et al., 2017).Pagel et al. (2007), for example, found evidence for the inverse correlation between frequency of use and rate of change across different parts of speech in four different Indo-European languages (English, Greek, Spanish, and Russian) but could only speculate on what gives rise to this pattern. Similarly,Lieberman et al. (2007) found strong support for the existence of this pattern in the regularization of English past-tense verbs over the past 1,200 years but could not provide an explanation for the pattern. Likewise examining the regularization English past-tense verb, Newberry et al. (2017) were able to detect signatures of selection in some cases (e.g., wove → weaved) but not in others (e.g., spilt → spilled). More importantly, however, their study was not designed to determine whether the overall inverse correlation between frequency of use and rate of change is due to drift or selection.

Moreover, these studies were based on corpus data. Corpus studies deal with recorded data from natural languages, but they cannot easily track the entire trajectory of a language or control the many different factors that affect language change (cf.Galantucci et al., 2012). Furthermore, corpus-based methods for inferring drift and selection can be sensitive to choices of data binning: A test for selection versus a null hypothesis of neutral drift was shown to depend on whether a corpus is parsed into time intervals of equal amount of data or equal duration of time (Karjus et al., 2020). Different results were also obtained in a binary classification of drift versus selection when analyzing the same corpus with a deep neural network (Karsdorp et al., 2020).

A solution is to complement such corpus-based approaches with experimental studies that permit greater control of relevant factors and that eliminate questions of data binning. Artificial-language experiments in particular allow the entire trajectory of the language to be recorded (Roberts, 2017). Such experiments also make it possible to isolate different linguistic, social, and communicative factors that affect language learning and to control and manipulate them (Folia et al., 2010; Culbertson and Schuler, 2019; Roberts and Sneller, 2020; Kanwal et al., 2017). This allows the problem of potential confounds to be reduced and causal relationships to be identified more easily. Such experiments are thus a very important tool for understanding the role of frequency effects in language change.

We here report the first such experiment designed explicitly to quantify the role of drift and selection in the relationship between word frequency and regularization. The experiment focuses specifically on drift and selection in learning, which is widely considered to be an important driver of language change (Kroch, 2005; Lightfoot, 2010; Labov, 2011; Sneller et al., 2019; Ferdinand et al., 2019). Indeed, language-learning experiments have already revealed several factors—e.g., age, memory, and multigenerational transmission—that up- or down-regulate language change during acquisition (Kirby et al., 2015; Hudson Kam and Newport, 2005; Samara et al., 2017; Perfors, 2012; Ferdinand et al., 2019; Reali and Griffiths, 2009; Smith and Wonnacott, 2010). Similar experiments have also examined the role of drift and selection in the emergence of simple communication systems (Tamariz et al., 2014). But no experiment to date has investigated if it is drift or selection that drives the negative correlation between frequency of use and regularization.

To study this, we conducted an experiment in which participants were tasked with learning a miniature artificial language. The language consisted of two nouns and two plural markers. During language acquisition, participants encountered nouns that occurred at different frequencies and plural markers that were associated with nouns at different relative frequencies. Participants were therefore subjected to drift and selection of varying strengths. By measuring the regularization of plural marking in the language, we were then able to determine whether low-frequency nouns did in fact regularize more than high-frequency nouns and, if so, whether it was drift or selection that was responsible for the greater regularization of low-frequency nouns. Our experimental setup therefore allowed us to test three main hypotheses: First, that greater regularization of low-frequency words results from stronger drift on low-frequency words (Hypothesis 1); second, that greater regularization of low-frequency words results from stronger selection on high-frequency words (Hypothesis 2); and third, that greater regularization of low-frequency words results from both selection and drift (Hypothesis 3).

## Method

Our experiment was pre-registered with the Open Science Foundation (https://osf.io/72kqa/?view_only=d0f2776850fe4c5e970b68237049c400).

### Participants

We recruited 400 participants through Prolific. To be eligible, participants had to report English as a native language. Participants were informed that this was a study on an alien language and were asked to give their consent before taking part in the experiment. Participants were paid a base rate of $1.00 for participating in the study and told that they would receive a 50% bonus depending on the accuracy of their answers; in reality, all participants who completed the study were given the 50% bonus. Data from participants who were more than two standard deviations in either direction from the mean completion time were discarded. There were ten such participants.

### Alien Language

Participants were trained on an artificial language composed of nouns for two different referents embedded in an English sentence. To facilitate learning, each noun consisted of a root with two syllables. Each root syllable consisted of a consonant followed by a vowel, with each of the two consonants matching a consonant in the corresponding English word (“buko” for book and “hudo” for hand). For each root, participants were asked to learn a singular and a plural form. The singular form consisted in the unmarked root; the plural was formed by adding a suffixed marker to the root with two possible variants, “-fip” and “-tay”, following Smith and Wonnacott (2010). Nouns belonged to one of two frequency classes: the low-frequency noun was shown six times during the training and testing phases; the high-frequency noun was shown 18 times. Frequency classes were therefore comparable to the ones used Kanwal et al.‘s (2017) study of another large-scale regularity found by Zipf. For each participant, plural markers were randomly assigned to noun and nouns were randomly assigned to frequency class.

### Procedure

Participants interacted with a custom-made website programmed with PennController for Ibex (Zehr and Schwarz, 2018), an online experiment scripting tool, and hosted on the PCIbex Farm (expt.pcibex.net). Instructions were provided on screen before each stage of the experiment. The experiment began with a training phase in which participants were asked to learn an alien language; we call the language that participants learned the *input language.* The training phase was followed by a testing phase in which participants were asked to use the language; we call the language that participants produced the *output language.*Participants passed through the following phases:

1. Training phase
  a. Noun Training Participants were shown a picture depicting a single object (Figure 1). Below the image, a caption with the sentence “Here is one buko” or “Here is one hudo” instructed participants on how to use the singular nouns. After clicking a *Next* button, participants were shown an image depicting another object. Each picture was shown once in random order, with a 300 ms pause between trials. Participants were then shown the same pictures two more times, alternating between a trial in which they were shown an object with the corresponding caption and a trial in which they were shown an object and asked to complete a sentence of the form “Here is one“. Participants had to enter the correct form of the noun to move on to the next trial. If the form was correct, participants were told so; if the form was incorrect, a box popped up reminding them of the correct form and asking them to try again.
  b. Plural Training Participants were shown a picture depicting three instances of the same object. The objects were the same as the ones shown during noun training. After clicking a *Next* button, participants were shown another image. Below each image, a caption with the sentence “Here are several buko+MARKER” or “Here are several
2. Testing phase Participants were shown pictures depicting three instances of the same object and with the same frequency as in the plural training phase. At random intervals, participants were shown the image of a singular object; singular objects were shown only once. Participants were asked to complete the sentence in each trial and therefore had to enter either the singular or plural form of the noun, depending on the picture shown. In the plural case, participants were told that the form was correct provided that it was seven characters long and that it contained one of the two plural markers at the end. If it was incorrect, participants were asked to try again without being told what the correct form was. In the case of the singular, participants were told that the form was correct provided that their answer was four characters long. Otherwise, participants were asked to try again without being told what the correct form was.

**Figure 1:**
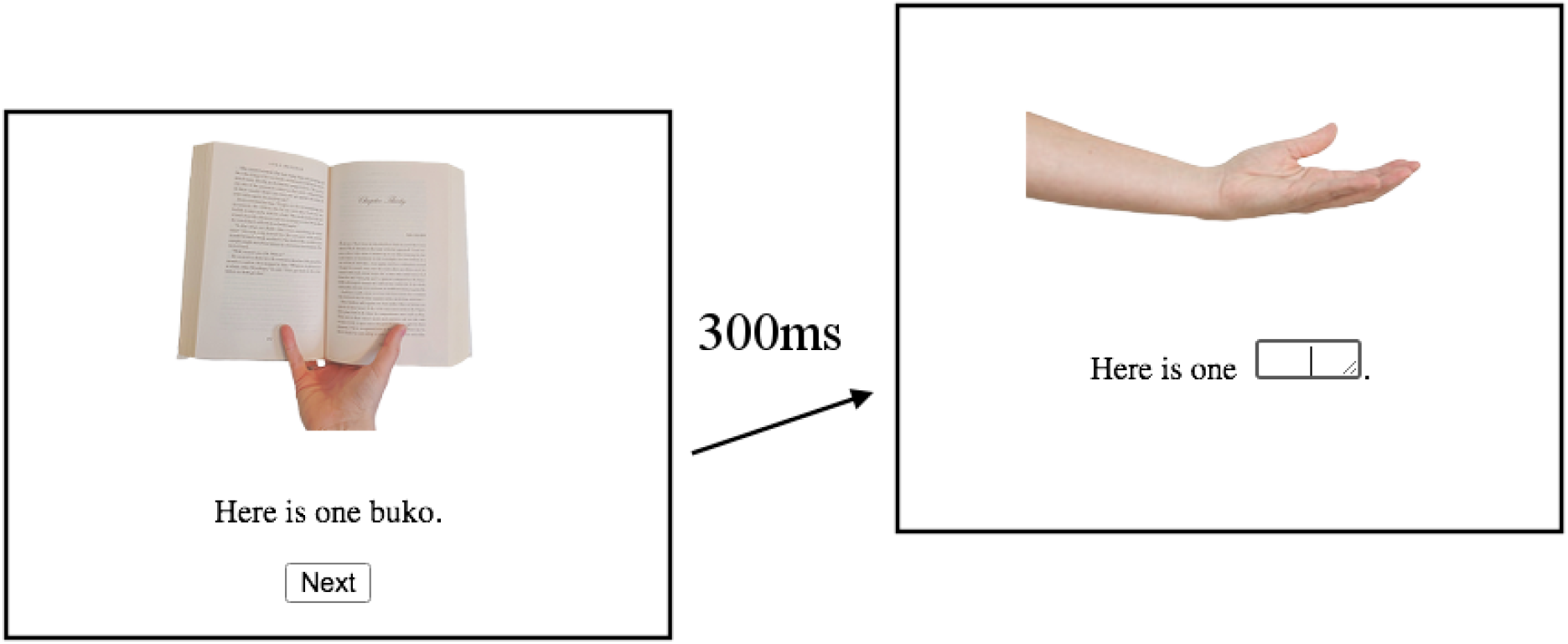
Training and Testing during Noun Training. hudo+MARKER” (where MARKER was either “fip” or “tay”) instructed participants on how to use the plural nouns. There was variation in how markers were associated with nouns (see *Conditions* Section). Depending on frequency class, each picture was shown either six or 18 times. Pictures were randomly selected to appear in each trial, with a 300 ms pause between trials. At random intervals, participants were shown the image of a singular object and asked to complete the sentence with the correct noun; this was done for each singular object only once. If the form entered was incorrect, a box reminded participants of the correct form and asked them to try again.

### Conditions

As discussed above in the *Alien language* Section, we manipulated noun frequency as a within-subjects variable such that one noun (the low-frequency noun) was shown to participants six times in training and the other (the high-frequency noun) was shown eighteen times.

We also manipulated the presence of selection as a between-subjects variable by manipulating the relative frequency of plural markers. In the *Drift Condition*, plural markers in the input language occurred at the same relative frequency during plural training: The ratio of plural markers associated with each noun was 1:1 (see Figure 2, *left*). The low-frequency noun therefore occurred with one marker in three trials and with the other marker in the remaining three trials; the high-frequency noun occurred with one marker in nine trials and with the other marker in the remaining nine trials. The purpose of the Drift condition was to establish an input language in which there was no directional pressure for regularization due to relative frequency, as randomization ensured that participants had no stimulus-related reasons for a bias in learning one or the other marker. If the language changed, it would be as a result of drift rather than selection.

**Figure 2:**
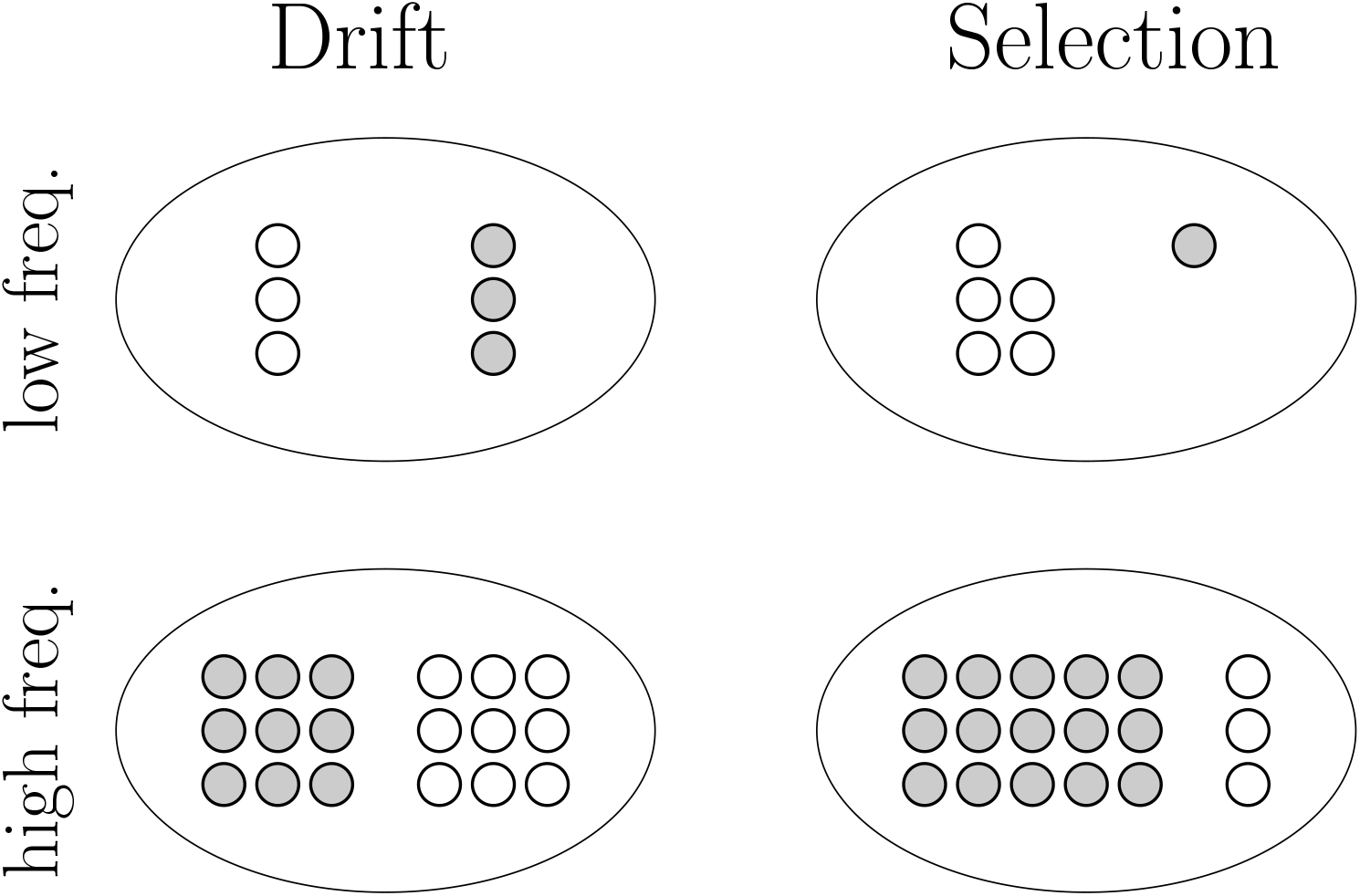
Drift & Selection Conditions. Each circle represents a noun in the input language; colors (white and gray) represent the two plural markers. In the Drift Condition, the ratio of plural markers associated with each noun was 1:1. In the Selection Condition, the ratio of plural markers associated with each noun was 5:1.

In the *Selection Condition,* plural markers in the input language occurred at different relative frequencies: the ratio of plural markers associated with each noun was 5:1 (see Figure 2, *right*). The low-frequency noun therefore occurred with one marker in five trials and with the other marker in the remaining trial; the high-frequency noun occurred with one marker in 15 trials and with the other marker in the remaining three trials. To facilitate learning, low- and high-frequency nouns differed with respect to which marker was more common. For example, if the low-frequency noun occurred five times with “-fip” and only once with “-tay”, then the high-frequency noun occurred 15 times with “-tay” and three times with “-fip”. The purpose of the Selection Condition was to establish an input language in which there was an asymmetry in the relative frequency of plural markers and thus the potential for a directional pressure—that is, selection—for one form at the expense of the other. In particular, we predicted that participants would adopt the more common marker with a probability greater than expected by chance alone.

A subtlety of design is worth mentioning here, as frequency was used both to manipulate the presence of selection and to manipulate the strength of drift. These two distinct uses of frequency in fact depended on the structure of the meaning space. In particular, the relative frequency of the nouns was not expected to be a significant source of selection because there was only one noun corresponding to each meaning, so there was no competition for meaning in the nouns. There was, however, competition between suffixes for indicating plurality, making the frequency bias a source of selection with respect to them.

### Dependent variable: regularization

For each noun, the more common marker in the Selection Condition was designated as the “primary” marker and the less common plural maker the “secondary” marker. For comparison, we arbitrarily labeled markers as “primary” or “secondary” in the Drift Condition as well. Following Lieberman et al. (2007), nouns in both conditions that occurred at least once with both markers were designated as the”irregular” nouns; nouns occurring with a single marker were termed “regular”. In the input language for both conditions, all nouns were irregular; in the output language, nouns could be either regular or irregular depending on the behavior of the participant.

To measure regularization, we calculated a *Regularization Index* for each participant (Lieberman et al., 2007). The Regularization Index (RI) was defined as the proportion of regular nouns in the output language. For each noun, the RI could therefore take a value of either 0 (for an irregular noun) or 1 (for a regular noun), such that a language with two regular nouns would have an RI of 1, and a language with one regular and one irregular noun would have an RI of 0.5. RI values were validated on the basis of conditional entropy, another commonly used measure of regularization (see Supplementary Material A).

### Pilot experiment

To test the viability of our experimental design, we conducted a pre-registered pilot experiment using the method described so far on a different sample of 400 participants (https://osf.io/ryc3j/?view_only=0026aa2c67184e339d2d7d478a8024ac). We report the results of this experiment in Supplementary Material B. The results revealed heterogeneity in the participant pool with respect to the experimental task. In the Drift Condition, the distribution of marker counts was trimodal: Most participants randomized their choice of markers, but some chose one or other of the markers exclusively (Supplementary Material, Figure 3 *left*). In the Selection Condition, the distribution of marker counts had a single peak and a long tail: While most participants chose the secondary marker with probability equal to or less than its initial frequency, many randomized their choice of markers (Supplementary Material, Figure 3 *right*). Results from the pilot experiment informed the statistical analysis plan for our main study, which is reported below.

**Figure 3:**
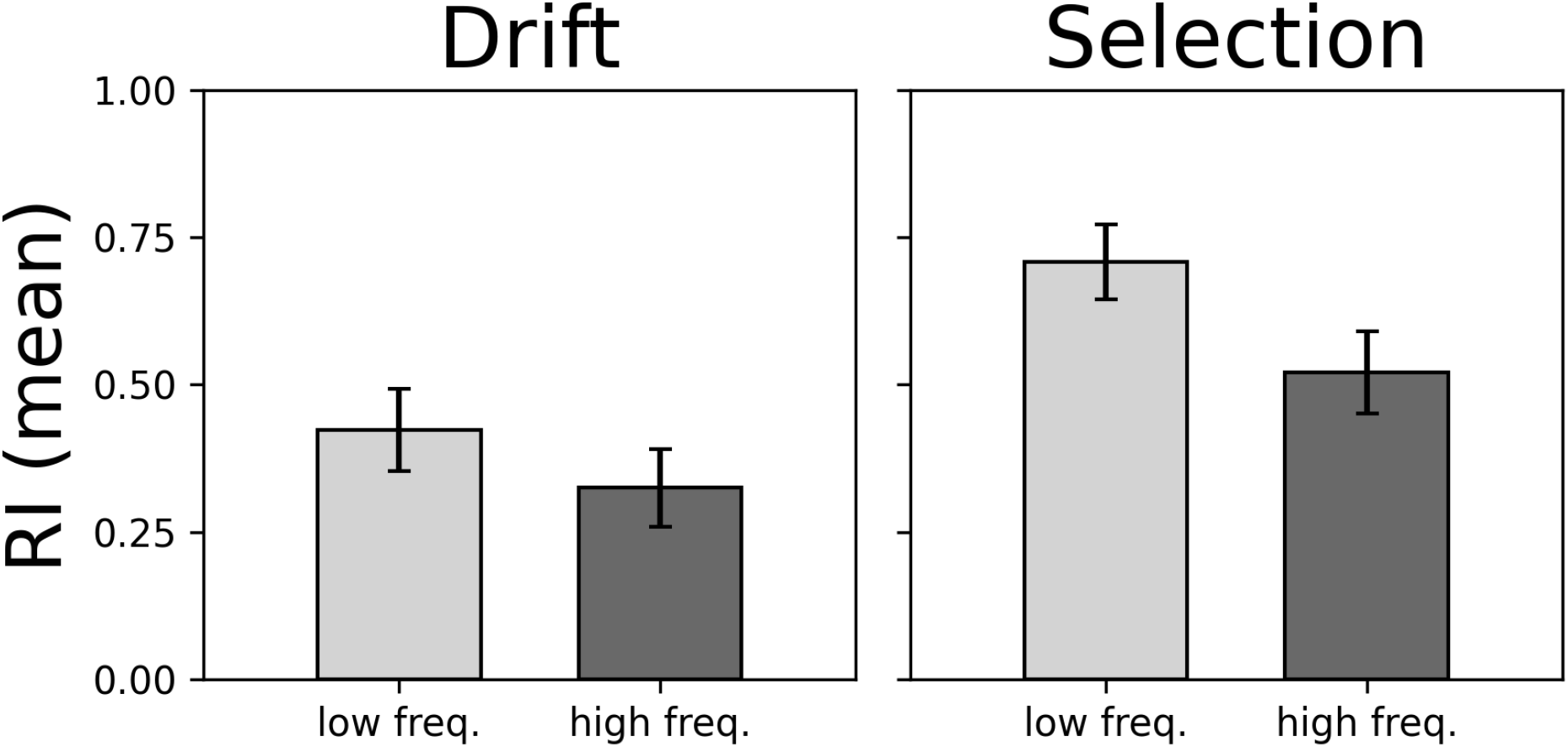
Regularization Index (RI). The mean RI is the proportion of regular nouns in the output language (error bars show 95% confidence interval). Drift: *N* = 194. Selection: *N* = 196

### Statistical Analysis

#### Distinguishing hypotheses

To distinguish between the three hypotheses, we used a binomial logistic model with noun regularity (i.e., regular or irregular) as the dependent dichotomous variable and frequency (i.e., low or high frequency) and selection (i.e., presence or absence) as independent dichotomous variables. In particular, the model took the following form:

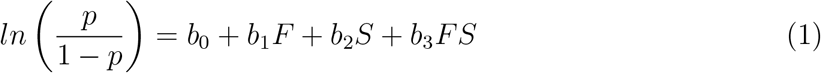

where *p* is the proportion of regular nouns, *F* indicates frequency class (low: 0; high: 1), *S* indicates absence or presence of selection depending on condition (absence of selection, ie. Drift Condition: 0; presence of selection, ie. Selection Condition: 1), and FS represents the interaction between frequency class (*F*) and selection (*S*). In this model, *b*_1_ measures the main effect of frequency class, *b*_2_ measures the main effect of selection, and *b*_3_ measures the interaction of frequency class and selection on noun regularization. Hence, if *b*_1_ differs from zero but *b*_3_ does not, the model supports the hypothesis that low-frequency forms regularize more because of drift (Hypothesis 1); if *b*_3_ differs from zero but *b*_1_ does not, the model supports the hypothesis that low-frequency forms regularize more because of selection (Hypothesis 2); and if both *b*_1_ and *b*_3_ differ from zero, the model supports the hypothesis that low-frequency forms regularize more because of a combination of drift and selection (Hypothesis 3). If *b*_2_ differs from zero, this does not correspond to any of the hypotheses we test but it conforms to the assumption that forms would regularize more overall in the Selection than in the Drift Condition.

#### Manipulation check

We used inferences under a Wright-Fisher model as a manipulation check to confirm the presence of selection for the primary marker in the Selection Condition. The Wright-Fisher model, commonly used in evolutionary biology and shown to be equivalent to models of iterated learning (Reali and Griffiths, 2010), describes a *population* of constant size *n* with discrete types and discrete generations. In our experiment the population in question was the ensemble of markers in a given frequency class (note that “population” here refers to the population of linguistic entities and not the population of language users). With two types (A and B), the probability that a population with *i* markers of type A and *n* – *i* markers of type B transitions to the next generation with *k* markers of type A and *n* – *k* markers of type B is given by a binomial distribution with parameters *n* and *f* (*n*, *s*), where *f* (*n*, *s*) is the success probability. The success probability is a function of *s,* the selection coefficient that measures the strength of selection for or against each one of the markers.

In our experiment, a population in the Wright-Fisher model corresponds to the ensemble of plural markers in a given frequency class. Accordingly, the population size *n* takes two different values depending on the frequency class: *n* = 6 in the low-frequency class, and *n* = 18 in the high-frequency class. Marker tokens correspond to individuals and marker types correspond to types in the Wright-Fisher model. In the Selection Condition, we treat the secondary marker as the focal type independently of the particular form that the marker may take (i.e., “-fip” or “-tay”). In the Drift Condition, we assign the labels “primary” and “secondary” to plural markers arbitrarily but in equal proportion to allow for comparisons between conditions. In both conditions, the input and output languages correspond to two distinct generations of the Wright-Fisher population.

As our pilot study revealed a heterogeneous participant pool, we first built a model representing different types of participant. The model assigned probability *q* that participants choose a single marker regardless of input language (“full regularizers”). Further, the model assigned probability *r* that participants choose markers according to a binomial distribution with parameters *n* and 0.5, where n is the number of trials in which a given noun appears and 0.5 means that participants randomize their choice of markers (“randomizers”). Finally, the model assigned probability 1 – *q* – *r* that participants choose markers according to the Wright-Fisher model with selection (“partial regularizers”).

As our population, for the purposes of the Wright-Fisher model, was defined as the ensemble of markers for a given frequency class, the population size *n* took different values depending on frequency class (*n* = 6 or *n* = 18). The success probability was given by:

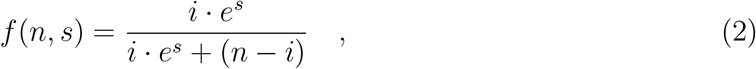

where *i* is the marker count for a given frequency class, *n* indicates the frequency class, and *s* is the selection coefficient for the secondary marker (i.e., *s* is positive if participants favor the secondary marker, negative if participants favor the primary marker, and exactly zero if participants show no preference for one marker or the other); we take the constant *e* to the power of *s* to ensure that *f*(*n*, *s*) is symmetrical about zero.

To estimate *s* using the Wright-Fisher model, we computed the likelihood of transitions from the initial state of the population (i.e., the input language) to the final state of the population (i.e., the output language) given different values of *s*. The maximum-likelihood estimate of selection was then the value of the selection coefficient that maximized the sum of the log-likelihood of transitions observed across participants. In our experiment, we estimated *s* together with the composition of the population. The estimate *ŝ* was therefore the value that maximized the sum of the log-likelihoods together with the proportions of randomizers 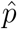 and full regularizers 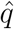. In particular, *ŝ* was given by the following expression:

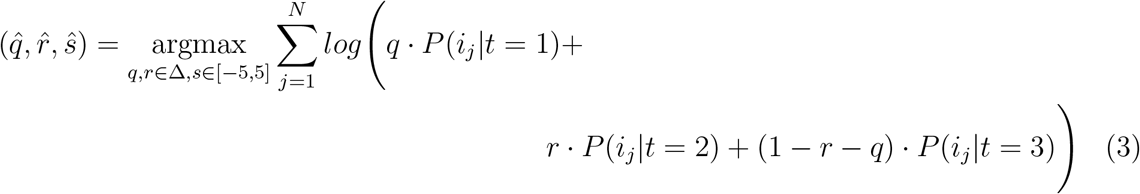

where *P*(*i_j_* |*t* = *k*) is the probability of participant *j* producing an output language with *i* secondary markers given that the participant is of type *k*, *t* = 1 if participant *j* is a full regularizers, *t* = 2 if participant *j* is a randomizer, and *t* = 3 if participant *j* is a partial regularizer. Here, Δ denotes the simplex volume {(*q*, *r*) ∈ [0,1]^2^ | *q* + *r* ≤ 1}. In this way, we simultaneously obtained *ŝ* among partial regularizers and the composition 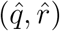 of the participant pool; we limit the estimate of *s* to the interval between (−5,5) as this included both the point estimate and confidence intervals for s in the analysis shown below.

The two-tailed 95% confidence interval for *ŝ* was given by the log-likelihood ratio and thus included values of *s* satisfying *ℓ*(*s*) – *ℓ*(*ŝ*) ≤ 1.92, where *ℓ*(*s*) is the sum of log-likelihood given *s* maximizing over parameters (*q*, *r*). Similarly, the two-tailed 95% confidence regions for 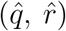 included all values of (*q*, *r*) satisfying 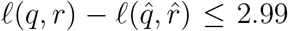, where *ℓ*(*q*, *r*) is the sum of log-likelihood given (*q*, *r*), maximizing over the parameter *s*.

Analysis was conducted using Python (Van Rossum and Drake Jr, 1995) and Julia (Bezanson et al., 2017). Data and scripts for the experiment are available at https://osf.io/5m9ak/?view_only=aaa9660774964e76838903533f6f0c16.

## Results

The rate of regularization was higher for low-frequency nouns in both conditions (Figure 3). In particular, RI estimates for low- and high-frequency nouns were 0.42±0.07 and 0.32±0.07 respectively in the Drift Condition (*N* = 194) and 0.71 ±0.06 and 0.52±0.07 in the Selection Condition (*N* = 196). This conforms to the Zipfian pattern of inverse association between frequency of use and regularization.

To test whether this pattern was statistically significant and to help identify what was driving it, we used a binomial logistic regression model. The model revealed a negative effect of frequency class on noun regularity, with low-frequency nouns being significantly more likely to regularize across conditions (*b*_1_ = −0.42±0.21; *p* = 0.047; Table 1). As expected, selection had a large positive effect on noun regularity (*b*_2_ = 1.2 ±0.21; *p* < 0.0001). There was no interaction between frequency class and selection (*b*_3_ = −0.39 ±0.29; *p* = 0.19). Given that frequency class had a significant effect on regularization but the interaction between frequency class and selection did not, our results provide support for the hypothesis that the greater regularization of low-frequency nouns was due to drift rather than selection.

**Table 1:** Logit model. Model was given by 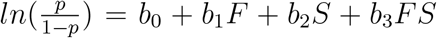. Significant results at the 0.05 level are marked with ‘*’.

| | $\beta$ | SE | $p$ |
| --- | --- | --- | --- |
| intercept ($b_0$) | -0.31 | 0.14 | 0.032* |
| frequency ($b_1$) | -0.42 | 0.21 | 0.047* |
| selection ($b_2$) | 1.2 | 0.21 | < 0.0001* |
| freq. × sel. ($b_3$) | -0.39 | 0.29 | 0.19 |

However, this conclusion only holds if selection was in fact present in our experiment. We therefore conducted a manipulation check (estimating selection under a Wright-Fisher model), to ensure that there was in fact selection against the secondary marker in the Selection Condition and no selection in the Drift Condition. The distribution of marker counts was similar to that obtained in our pilot study: in the Selection Condition, the distribution of marker counts had a single peak and a long tail; in the Drift Condition, the distribution of marker counts was trimodal (Figure 4). We therefore sought to determine whether there was selection in the Selection Condition and no selection in the Drift Condition using our population model.

**Figure 4:**
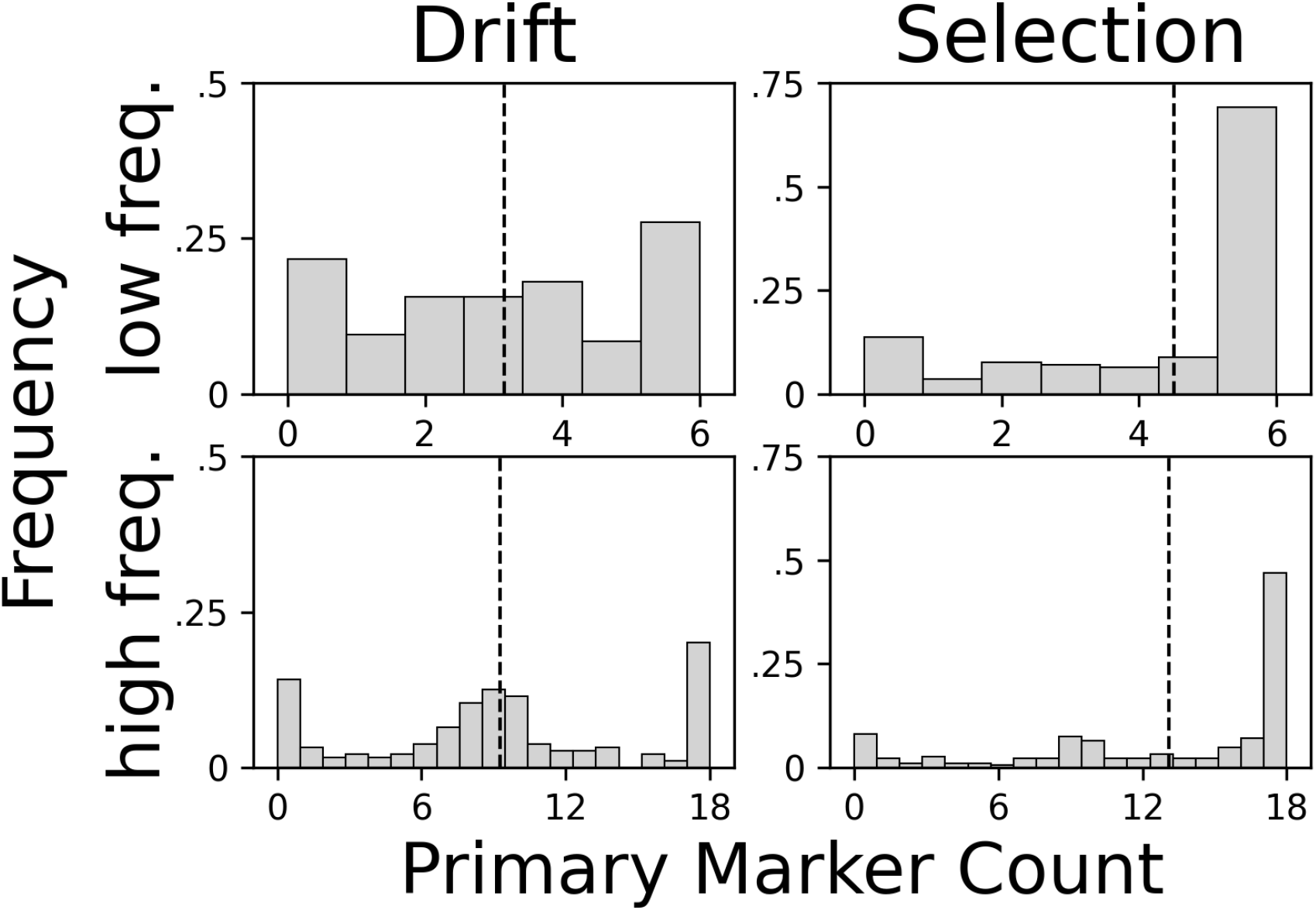
Distribution of primary marker counts. Empirical distribution shown by gray bars; mean shown by dashed line. In the Drift Condition, the distribution was trimodal. In the Selection Condition, the distribution had a single peak with a long tail.

In the Selection Condition we found evidence of selection against the secondary marker: Among partial regularizers, *ŝ* was equal to −2.3±(0.9, 0.6) and −2.1 ± (0.3, 0.4) for low- and high-frequency nouns (Figure 5). Estimates for low- and high-frequency nouns were similar in value, indicating that selection was of comparable strength in both frequency classes.

**Figure 5:**
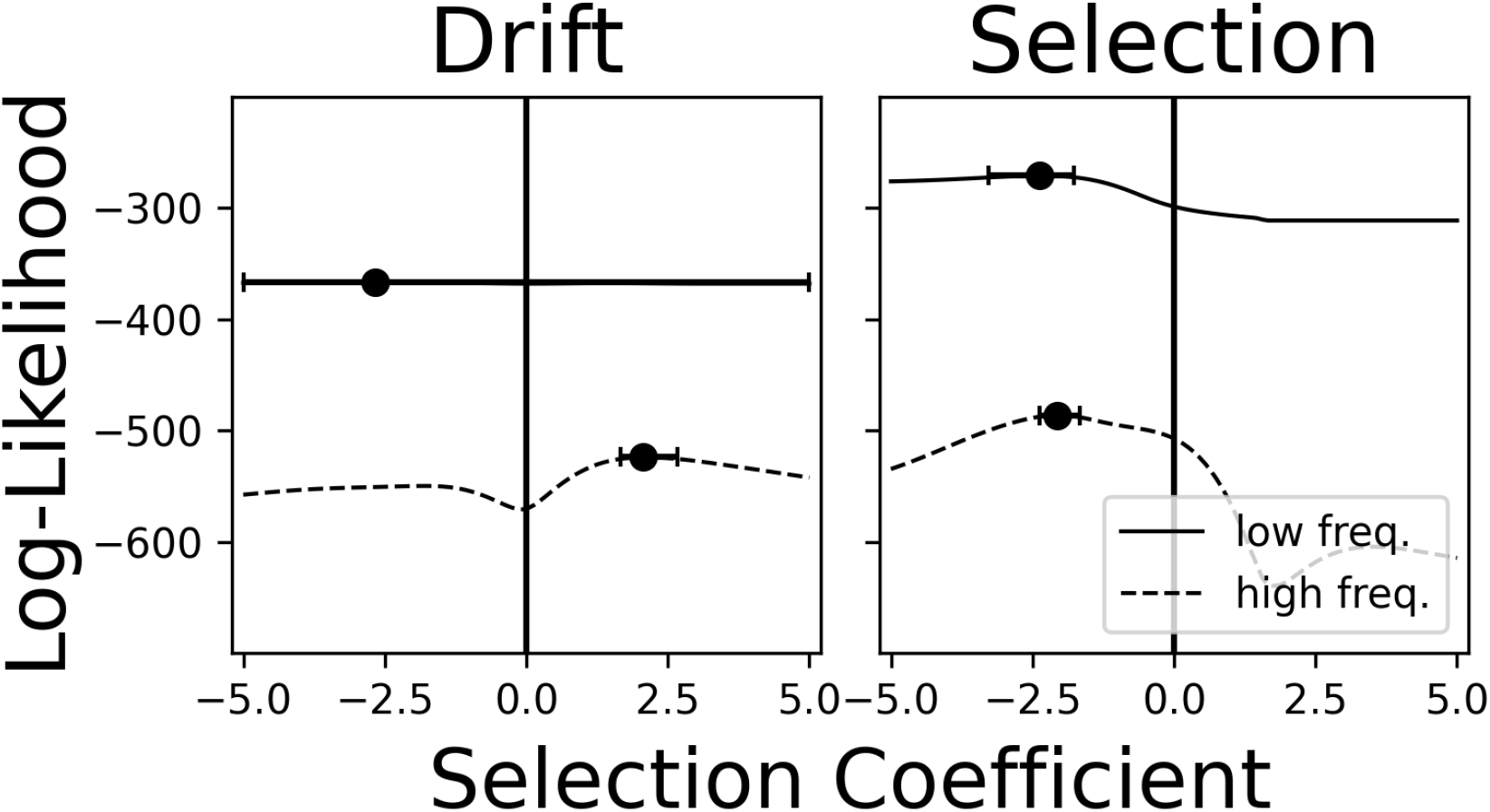
Lines indicate the sum of log-likelihoods for the data given the selection coefficient for partial regularizers in the population model. Circles show maximum-likelihood estimates of the selection coefficient (i.e., the value of the selection coefficient that maximizes the likelihood of the observed data); error bars show two-tailed 95% confidence intervals.

In the Drift Condition, there was no evidence of selection among partial regularizers in the low-frequency class: *ŝ* was −2.1±(2.93, 7.1), with the 95% confidence interval spanning the entire range of selection coefficients sampled (Figure 5). Contrary to our expectation, however, our estimate for the selection coefficient was positive in the high-frequency class: *ŝ* was 1.97 ± (0.4, 0.71). This might seem like an indication that there was selection in the Drift Condition, which would clash with a central assumption of our analysis plan. But this was likely not the case. In the Drift Condition, estimates for the proportion of randomizers in the low- and high-frequency classes were 0.56 and 0.6 (Figure 6). Similarly, estimates for the proportion of full regularizers were 0.37 and 0.31. Both participant types therefore made up almost the entirety of the sample, with partial regularizers comprising only 0.07 and 0.09 in the low- and high-frequency classes. It is thus likely that our maximum-likelihood algorithm detected positive selection among partial regularizers in the high-frequency class due to the small number of partial regularizers in the sample: We estimated that there were very few partial regularizers in our sample (namely, 0.09 × 193 ≈ 17), and the chi-squared asymptotic confidence interval on maximum-likelihood estimates is a poor approximation in small samples. Negative selection was detected in our pilot study, corroborating this point (see Supplementary Material B, Figure 4).

**Figure 6:**
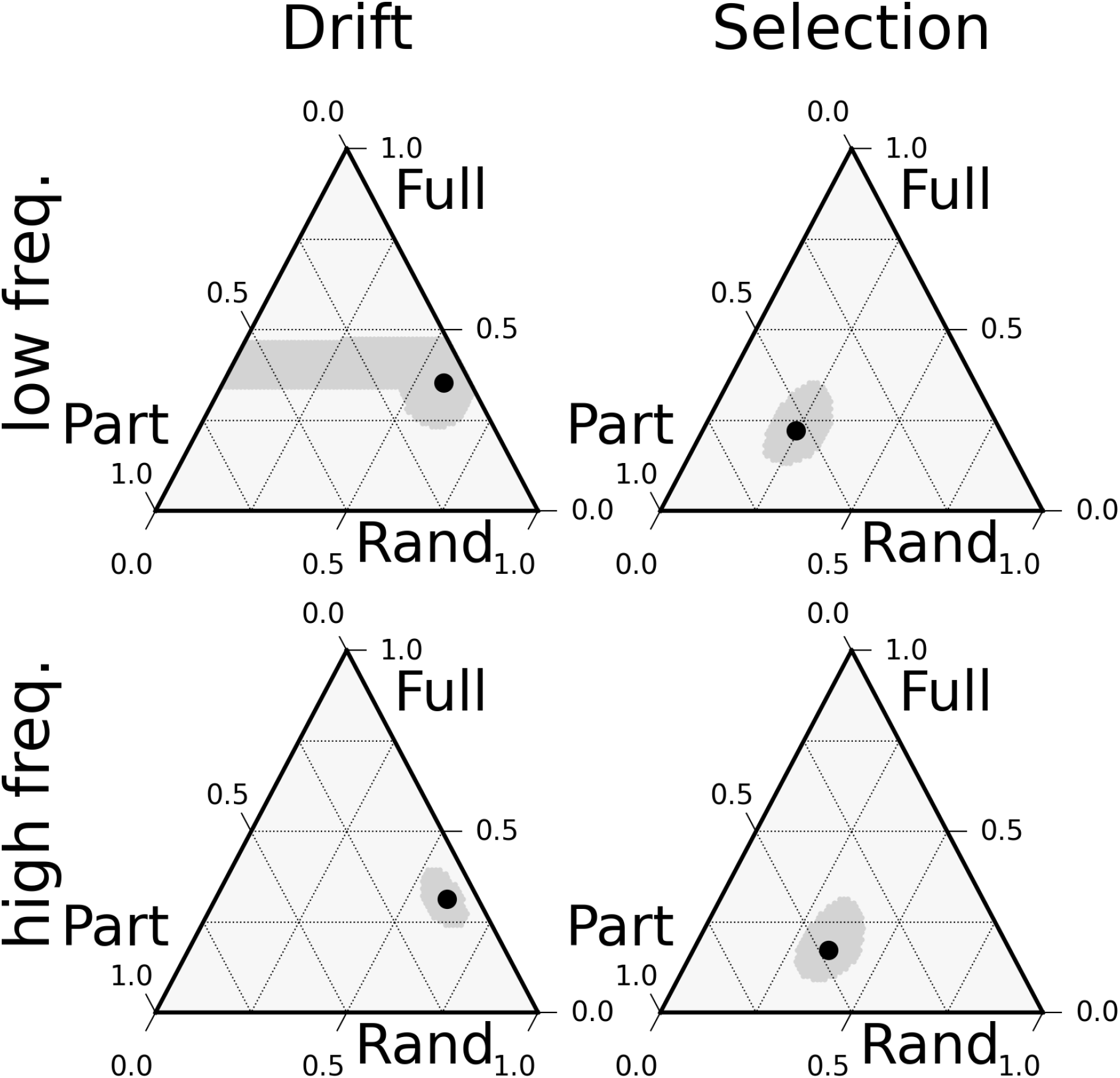
Population Composition. Black circles show the maximum-likelihood composition of the population with proportion *p* of randomizers, proportion *q* of full regularizers, and proportion 1 – *p* – *q* of partial regularizers; 95% confidence regions shown in gray.

In the Selection Condition, we estimated that partial regularizers made up 0.54 and 0.48 of the population in the low- and high-frequency classes; the proportion of randomizers was 0.24 and 0.35, and the proportion of full regularizers was 0.2 and 0.17. Together with the finding that selection was negative in the Selection Condition, this therefore suggests that selection against the secondary marker was indeed present in the Selection Condition but likely absent in the Drift Condition.

Since the number of partial regularizers was higher among low-frequency nouns in the Selection Condition (0.54 vs. 0.48), and drift should be stronger at low frequencies, these results provide further support for the hypothesis that low-frequency nouns regularized more because of drift. Moreover, selection among partial regularizers was approximately equal for low- and high-frequency nouns in the Selection Condition: *s* = −2.3 ± (0.9,0.6) vs. *s* = −2.1 ± (0.3, 0.4). This means that the difference in regularization between low- and high-frequency nouns could not be due to a difference in selection. These results therefore support the hypothesis that drift alone was responsible for the difference in regularization. This is consistent with findings based on a comparison between RI across frequency classes in both conditions and our regression model.

This analysis yielded consistent results when applied to our pilot data (see Supplementary Material B).

## Discussion

We conducted an experiment in which participants learned a miniature language with irregular plural marking, and we manipulated the strength of drift and selection acting on the markers. We found a difference in regularization between low- and high-frequency nouns regardless of selection strength, indicating that the difference between frequency classes in our experiment was sufficient for the Zipfian pattern to emerge. The absence of an interaction between frequency class and selection suggests that this difference was not due to selection. Indeed, results suggest that drift during language acquisition was sufficient for generating the negative correlation between word frequency and regularization rate that Zipf first noted in natural languages (Zipf, 1949). Our study therefore adds to a growing body of evidence suggesting that drift may be a major driver of language change, including patterns of regularization.

Although this was not the primary goal of our study, our results also highlight the risk of assuming—rather than showing—that participants approach an experimental task as a homogeneous population. By exploring this in our own data, we were able to identify that our participant pool was not in fact homogeneous: in the Selection Condition, most participants regularized the use of plural markers, but many opted instead to randomize their choice of markers or to simplify the task by using a single marker throughout; in the Drift Condition, most participants randomized their choice of markers but many also simplified the task by using a single marker. Our experiment was not designed to determine why participants adopted such disparate strategies, but the results suggest that future work should account for potential heterogeneity in learning style, which is in part but not entirely influenced by the nature of the data (Siegelman et al., 2017; Hudson Kam and Newport, 2009). There may also be implications for language change that arise from the observed variability in learning styles, which remains a topic for future study.

There are several important limitations to our study. First, our experiment focused on only one source of selection (relative frequency). While we consider that this was a good place to start—in particular because it provides a purely frequency-based explanation for Zipf’s observation—it is not the only potential factor driving selection, so it remains possible that other sources of selection might play a role in the greater regularization of low-frequency nouns in natural language. For instance, selection might be stronger for high-frequency nouns if these words function as “anchors” during language acquisition and learning (Frost et al., 2019). It is also possible that factors such as morpheme length, phonological complexity, or iconicity might interact with frequency as sources of selection, further complicating the picture. Nonetheless, even if some other source of selection plays a role in producing the observed effect, our results suggest that drift is sufficient to produce the effect on its own.

Along similar lines, it is also worth noting that our study focused on the influence of drift and selection during learning and production. Interaction with other language users, which we did not incorporate into our task, might provide further sources of selection, such as selection related to social meaning and identity (Roberts and Fedzechkina, 2018; Sneller and Roberts, 2018) or communicative pressures (Galantucci, 2009; Wade and Roberts, 2020). While our results suggest that drift in language learning is a sufficient mechanism for generating the greater regularization of low-frequency terms, its role may be modulated by different forms of selection under certain circumstances. This would be an interesting focus for future work.

It is likewise important to consider the size of the artificial language. Consisting of two nouns and two affixes, it was the smallest possible language for our purposes. This was done to maintain careful control over how the language was learned: Participants were likely to learn the language in full rather easily, preventing differences in learning success from constituting a nuisance variable. It also made the experiment short and quick to run, which allowed us to gather a large sample cheaply and efficiently. A negative consequence, however, is that the ease of the learning task might have increased the potential for demand characteristics to play a role. We consider, however, that the downside of a simple language was outweighed by its benefits, since tight control over marker and noun frequencies was important.

In conclusion, we conducted the first (to our knowledge) experimental investigation of the role of drift and selection in explaining Zipf’s observation about word frequency and rate of change. We found that drift was sufficient to explain the pattern and that frequency-based selection did not play an important role. As this was an experimental study, we were able to control for and rule out other factors that make it difficult to discern the effect of drift and selection in corpus-based studies of natural languages—such as the effect of age of word acquisition, phonology, etc. Future work might expand the paradigm by employing more complex languages or incorporating more complex social contexts, including direct communication between participants (Wade and Roberts, 2020; Sneller and Roberts, 2018) or simulated communication (Buz et al., 2016). While our results suggest that drift in learning is sufficient to produce the observed relationship between word frequency and regularization, other language-internal and -external factors might play an important role as well. Future work could therefore investigate the role of selection of different strengths, or compare different sources of selection (Tamariz et al., 2014). There are also different possibilities for how regularization is operationalized. Following Hudson Kam and Newport (2005), Lieberman et al. (2007),Newberry et al. (2017),Ferdinand et al. (2019) and others, we operationalized regularization as the loss of competing forms for the same lexical item; it could, however, be operationalized differently, such as in terms of the loss of competing forms for the same meaning in the system as a whole. It might be that selection plays a more important role in such cases. The study we have presented here offers a simple and easily replicable paradigm for investigation of these and many related questions.

