## Supplementary Materials for "Drift as a driver of language change: An artificial language experiment"

### Drift as a driver of language change: An artificial language experiment: Supplementary Material

#### A Conditional entropy as a measure of regularization

A commonly used measure of regularization is the mean percentage change in the conditional entropy of linguistic forms (Ferdinand et al., 2019). Although this was not included in our analysis plan, we measured regularization as the mean percentage change in conditional entropy of marker on noun between input and output language to validate our results based on RI. The conditional entropy of marker on noun is defined as follows:

$$H(P|Q) = -\sum_{q_i \in Q} P(q_i) \sum_{P_j \in M} P(P_j|Q_i) \cdot log_2 P(P_j|Q_i) \quad (1)$$

where $P$ is the set of plural markers, $Q$ is the set of nouns, and $P(\cdot)$ is the probability with which participants use a linguistic form. Marginal and conditional probabilities were estimated as relative frequencies. The mean percentage change in conditional entropy between the input and output language can then be used as a measure of regularization. Higher positive values mean that the language gained entropy, i.e. the output language is less regular than the input language. Lower negative values mean that the language lost entropy, i.e. the output language is more regular than the input language.

Measured as mean percentage change in the conditional entropy of markers on nouns, regularization was higher for low-frequency nouns than high-frequency nouns in both conditions (Figure 1). In particular, the mean change in conditional entropy was $-0.5 \pm 0.06$ and $-0.40 \pm 0.06$ for low- and high-frequency nouns in the Drift Condition and $-0.62 \pm 0.08$ and $-0.45 \pm 0.09$ for low- and high-frequency nouns in the Selection Condition. These results are therefore consistent with the Regularity Index (RI), which also indicated higher regularization for low- than high-frequency nouns in both conditions.

#### B Results from Preliminary Study

Consistent with results from our main experiment, regularization was higher for low-frequency nouns in both conditions (Figure 2). In particular, RI estimates for low- and high-frequency nouns were $0.48 \pm 0.07$ and $0.33 \pm 0.07$ respectively in the Drift Condition ($N = 193$) and $0.71 \pm 0.06$ and $0.55 \pm 0.07$ in the Selection Condition ($N = 195$).

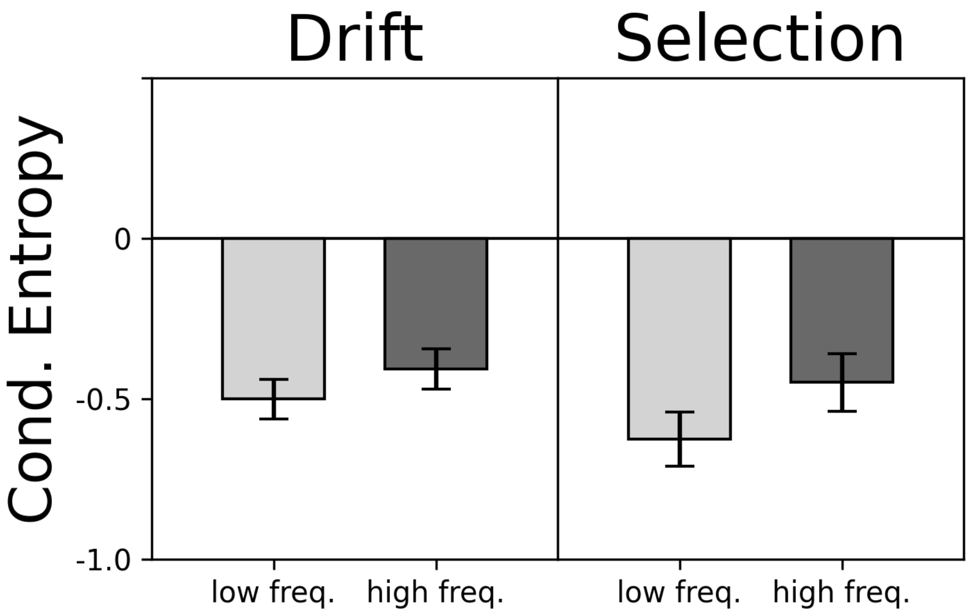


Figure 1: Change in conditional entropy of markers on nouns. Mean change in conditional entropy (bars) with 95% confidence intervals for the mean (error bars).

As in our main experiment, the distribution of marker counts in the Selection Condition had a single peak and a long tail; in the Drift Condition, the distribution of marker counts was trimodal (Figure 3). According to our manipulation check, estimates of selection among partial regularizers in the Drift Condition were again very low: $\hat{s}$ was equal to $-1.53 \pm (3.5, 6.5)$ and $-2.75 \pm (0.6, 0.8)$ in the low- and high-frequency classes (Figure 4). In the Selection Condition, estimates of selection for both frequency classes had roughly the same value: $\hat{s}$ was equal to $-2.55 \pm (0.8, 0.6)$ and $-2.55 \pm (0.6, 0.4)$ for low- and high-frequency nouns.

Estimates for the population composition were similarly comparable to those from our main experiment (Figure 5). In the Drift Condition, the proportion of randomizers was 0.5 and 0.62 for low- and high-frequency nouns, the proportion of full regularizers was equal to 0.44 and 0.30, and the proportion of partial regularizers was therefore 0.06 and 0.07. In the Selection Condition, the proportion of randomizers was equal to 0.24 and 0.35 for low- and high-frequency nouns, the proportion of full regularizers was 0.17 and 0.1, and the proportion of partial regularizers was 0.6 and 0.55.

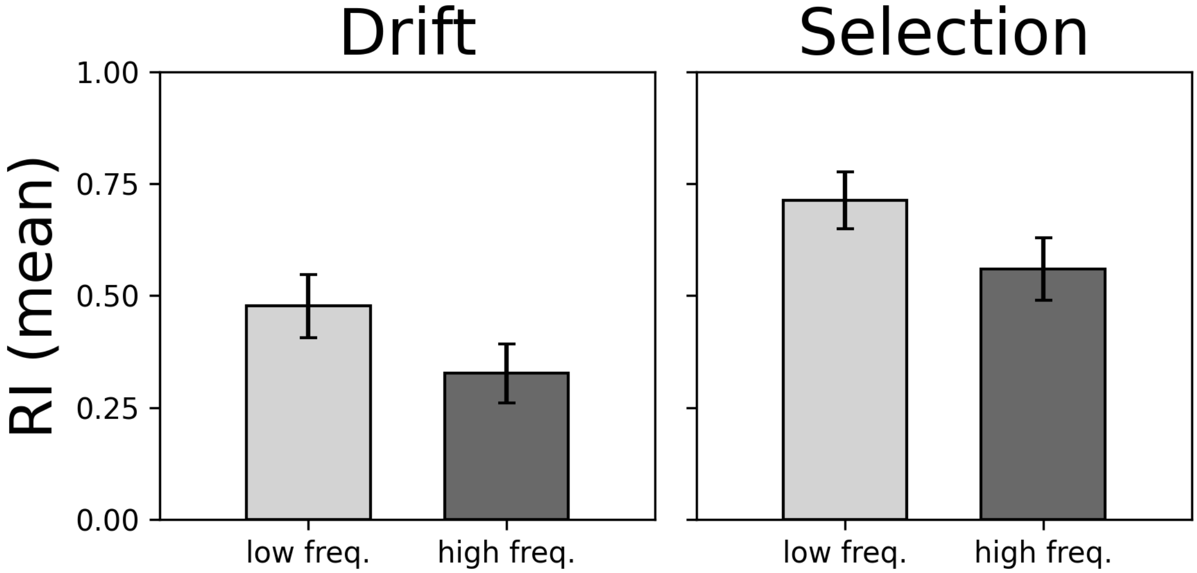


Figure 2: Regularization Index (RI). The mean RI is the proportion of regular nouns in the output language (error bars show 95% confidence interval). Drift: $N = 193$. Selection: $N = 195$

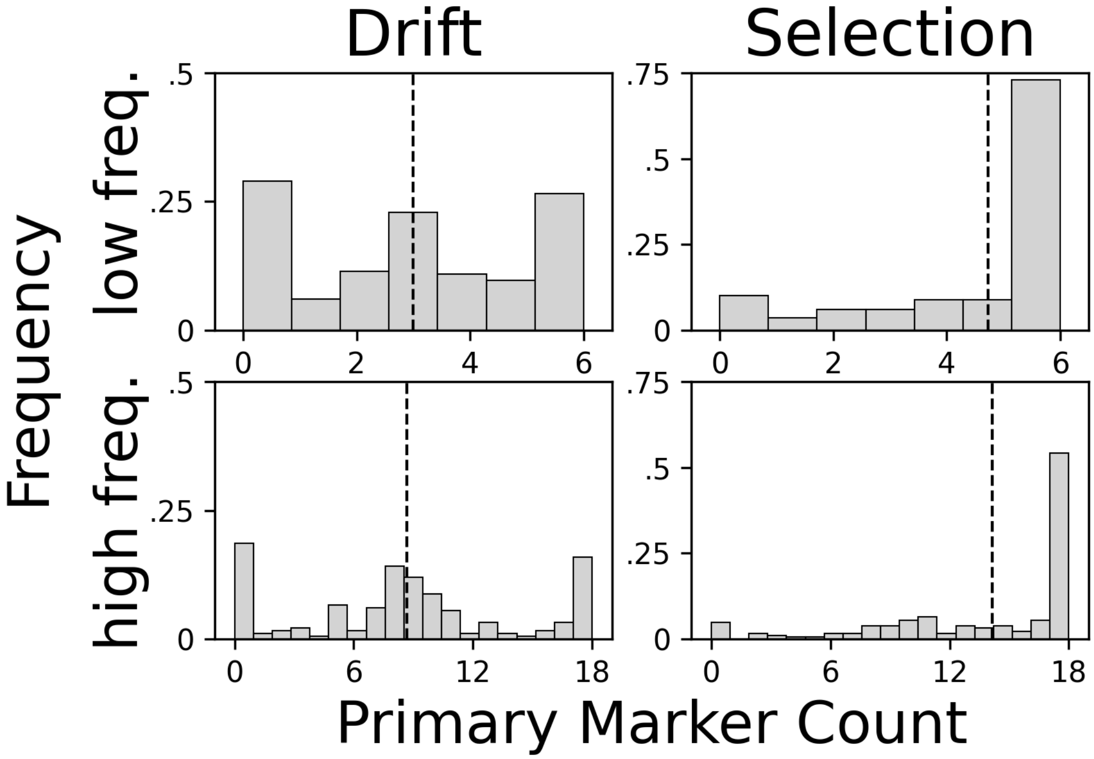


Figure 3: Distribution of primary marker counts. Empirical distribution shown by gray bars; mean shown by dashed line. In the Drift Condition, the distribution was trimodal. In the Selection Condition, the distribution had a single peak with a long tail.

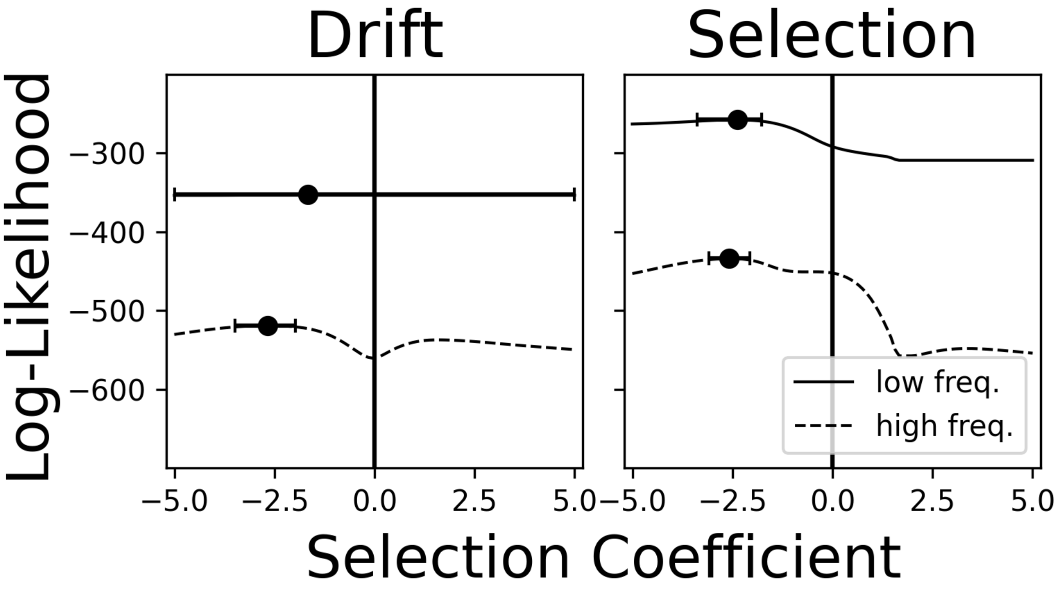


Figure 4: Lines indicate the sum of log-likelihoods for the data given the selection coefficient for partial regularizers in the population model. Circles show maximum-likelihood estimates of the selection coefficient (i.e., the value of the selection coefficient that maximizes the likelihood of the observed data); error bars show two-tailed 95% confidence intervals.

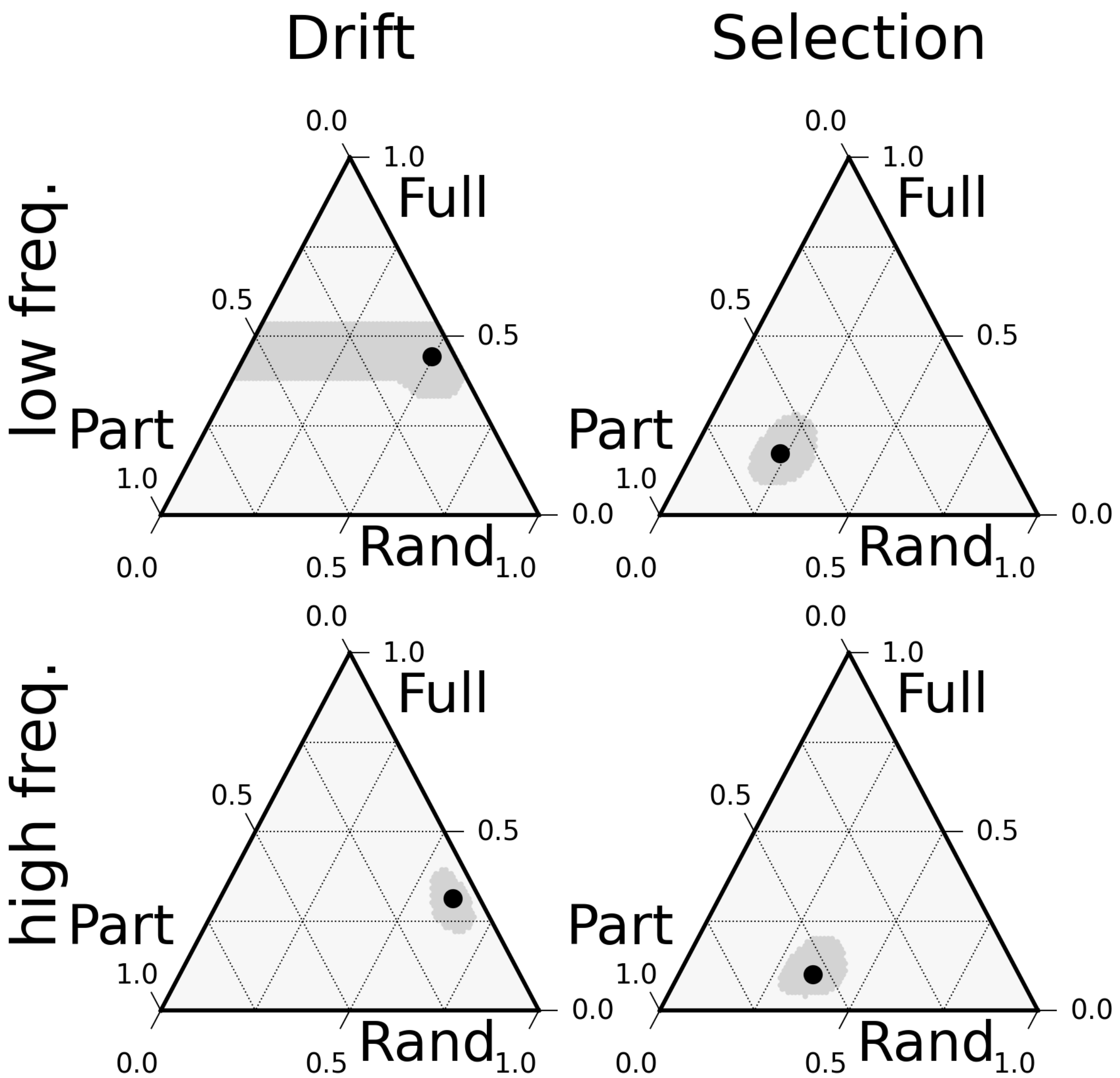


Figure 5: Population Composition. Black circles show the maximum-likelihood composition of the population with proportion $p$ of randomizers, proportion $q$ of full regularizers, and proportion $1-p-q$ of partial regularizers; 95% confidence regions shown in gray.
